# Modulation of Transferrin receptor by HIV-2

**DOI:** 10.1101/2025.07.03.663036

**Authors:** Aya Shamal Al-Muffti, Irene Wanjiru Kiarie, József Tőzsér, Mohamed Mahdi

## Abstract

Human immunodeficiency viruses (HIV-1 and HIV-2), are the causative agents of the acquired immunodeficiency syndrome (AIDS), that share substantial genomic and structural similarities, yet differ in replication dynamics and disease progression. While HIV-1 primarily enters host cells via CD4 and the chemokine co-receptors CCR5 and CXCR4, HIV-2 engages a broader range of chemokine receptors. The transferrin receptor (TFRC/CD71), a membrane protein essential for iron uptake and immune function, has recently been implicated in viral entry by other pathogens. In this study, we investigated the regulation of TFRC expression in cells transduced with HIV-1 and HIV-2 pseudovirions. We observed a significant upregulation of TFRC mRNA and protein levels in HEK-293T cells 27 hours post-transduction with HIV-2, but not with HIV-1. In Jurkat T-cells, TFRC transcript levels were significantly elevated at 36 and 40 hours following HIV-2 transduction. Mechanistic analyses revealed that this upregulation depends largely on a functional HIV-2 Tat protein. Additionally, HIV-2 transduction resulted in increased intracellular iron levels in Jurkat cells at 48 hours post-infection. Collectively, these findings identify TFRC as a novel Tat-dependent host target of HIV-2, shedding light on its broader receptor utilization and potential effects on host iron metabolism and viral tropism.

## Introduction

Human Immunodeficiency Viruses type 1 and type 2 (HIV-1 and HIV-2) are enveloped, single-stranded, positive-sense RNA viruses belonging to the family Retroviridae, subfamily Orthoretrovirinae, and classified under the Lentivirus genus. They are the causative agents of acquired immunodeficiency syndrome (AIDS) [1,2]. HIV-1, which is believed to have originated from the simian immunodeficiency virus in chimpanzees (SIVcpz) [3], is the most prevalent type worldwide [4,5]. In contrast, HIV-2 is thought to have originated from the simian immunodeficiency virus of sooty mangabeys (SIVsmm) and remains largely endemic to West Africa [6,7]. Although primarily confined to that region, earlier reports have documented sporadic increases in HIV-2 incidence in parts of Asia and Europe [7,8].

Compared to HIV-1, HIV-2 is characterized by lower transmissibility, reduced pathogenicity, and slower progression to AIDS [9,10]. Both HIV-1 and HIV-2 utilize the cluster of differentiation 4 (CD4) receptor for cellular entry, primarily targeting CD4+ T lymphocytes and other immune cells expressing this receptor [11,12]. Viral entry is also facilitated by coreceptors from the chemokine receptor family, predominantly C-C chemokine receptor type 5 (CCR5) and C-X-C chemokine receptor type 4 (CXCR-4) [12,13].

Upon encountering a target host cell, the HIV envelope surface glycoprotein (gp120 of HIV-1, and gp125 of HIV-2) engages the primary receptor CD4. A critical aspect of this interaction involves the insertion of the Phe43 residue of CD4 into a hydrophobic pocket on the viral glycoprotein, which induces structural rearrangements within the envelope. These conformational changes expose the binding site for a co-receptor (typically CCR5 or CXCR4) [1,14]. Thereafter, subsequent engagement of the co-receptor facilitates the activation of the fusion peptide located at the N-terminus of the transmembrane subunit (gp41 in HIV-1 and gp36 in HIV-2). This activation drives extensive conformational changes, ultimately resulting in the juxtaposition of viral and host membranes and the formation of a fusion pore [15].

To date, it remains a matter of debate whether fusion predominantly occurs at the plasma membrane or within endocytic vesicles [12,16,17].

HIV-1 entry is widely recognized as pH-independent, allowing fusion to occur without the need for acidic endosomal environments [18-20]. However, emerging evidence suggests that in certain cell types, such as HeLa-derived and CD4+ T cells, endocytosis is essential for complete fusion, involving dynamin and endosomal trafficking proteins like Ras-related protein Rab-5A (Rab5A) and sorting nexin family 3 (SNX3), despite the absence of pH-dependent mechanisms [21,22].

In contrast, some studies argue that plasma membrane fusion predominates in specific T-cell lines, highlighting cell-type-specific differences [18,23,24]. These findings underscore the complexity of HIV-1 entry pathways and the need for further research to clarify the contributions of endocytic and plasma membrane fusion [17].

The transferrin receptor 1 (TFRC) is a type 2 membrane protein with a molecular weight of 97 kDa, typically found as a homodimer on the cell surface of all vertebrates [25,26]. It is critical for cellular iron uptake, facilitating the endocytosis of iron-bound transferrin to deliver iron into cells for essential metabolic processes, such as hemoglobin synthesis and DNA replication [27,28], therefore, it is highly expressed on rapidly dividing cells, including erythroid precursors and cancer cells, due to their increased iron requirements [29]. In contrast, transferrin receptor 2 (TFRC2), while structurally similar to TFRC, exhibits a more restricted tissue distribution, being primarily expressed in the liver and erythroid progenitor cells. Unlike TFRC, TFRC2 displays a lower affinity for iron-bound transferrin, positioning it as a key sensor of systemic iron saturation levels [30].

In T cells, activation and proliferation trigger an upregulation of TFRC to enhance iron uptake, a process initiated by T cell receptor engagement with antigen–major histocompatibility complexes and increased IL-2 receptor expression [31,32]. The resulting intracellular iron surge is essential for T cell differentiation and clonal expansion. This transient TFRC elevation is regulated by a rapid endocytic sorting pathway, involving flotillin proteins and Rab5- and Rab11a-positive endosomes [33].

Several viruses were found to exploit TFRC as an entry point into host cells, notably arenaviruses [34], canine parvovirus, and feline panleukopenia virus [35], as well as certain strains of mouse mammary tumor virus (binding to mouse TFRC) [36], all binding to TFRC with high affinity to initiate infection. Given the high-level expression of TFRC on the cell membrane, it serves as a strategic entry route for pathogens via clathrin-mediated endocytosis. This viral exploitation of TFRC highlights its critical role in host-pathogen interactions and underscores its potential as a target for therapeutic intervention.

A recent study reported that Influenza A virus may also use TFRC recycling to enter host cells. [37]. Similarly, this receptor has been shown to bind the SARS-CoV-2 spike protein, thereby mediating viral entry [38].

In this study, we investigated the potential role of TFRC in the HIV life cycle, with a particular focus on its involvement in viral entry, considering the broad cellular tropism characteristic of HIV-2. To this end, we employed HIV-1 and HIV-2 pseudovirions pseudotyped with vesicular stomatitis virus glycoprotein (VSV-G). Our findings indicate that HIV-2, but not HIV-1, induces an upregulation of TFRC expression in the early phase of transduction.

## Materials and methods

### Plasmids and Vectors

HIV-1 and HIV-2 pseudoviruses were produced using second-generation lentiviral plasmids.

For HIV-1, the following constructs were used: packaging plasmid psPAX2 (a kind gift from Dr. Didier Trono at the University of Geneva Medical School, Geneva, Switzerland), transfer vector PWOX that was modified to code for mCherry [39], and pMD.G that encodes for vesicular stomatitis virus glycoprotein (Addgene, Watertown, MA, USA). For HIV-2, the plasmids consisted of HIV-2 CGP, a protein expression vector based on the ROD strain; HIV-2 CRU5SINCGW, which functions as the transfer plasmid encoding a GFP under a CMV promoter; and the pMD.G plasmid. HIV-2 plasmids were a kind gift from Joseph P. Dougherty from the Robert Wood Johnson Medical School, NJ, USA. A previously constructed HIV-2 CGP plasmid, which encodes a defective Tat protein containing the inactivating Y44A mutation, was also used in this study [40].

### Production of HIV-1 and HIV-2 pseudovirions

HEK-293T human embryonic kidney cells (Invitrogen, CA, USA) were maintained according to the manufacturer’s standard protocol. Cells were cultured in Dulbecco’s modified Eagle’s medium (DMEM) (Sigma-Aldrich, St. Louis, MO, USA), supplemented with 10 % Fetal Bovine Serum (FBS), 1 % L-Glutamine and 1 % penicillin-streptomycin. The day before transfection, cells in T-75 flask were passaged in order to achieve 60-70 % confluency the following day. On the day of transfection, the cells were transfected with the lentiviral vectors as described previously [41]. Briefly, a total of 30 μg plasmid DNA (HIV-2) or 36 μg plasmid DNA (HIV-1) were used for transfection, using a polyethylenimine (PEI) method (Sigma-Aldrich, St. Louis, MO, USA) [41]. 24 and 48 hours after transfection, the medium was collected and the virions were concentrated with the aid of Amicon Ultra-15 Centrifugal Filter Units (Sigma-Aldrich, St. Louis, Missouri, USA). The filtered virus was aliquoted into Eppendorf tubes and stored at –70 °C until further use. The activity of the produced pseudovirions was measured using enzyme-linked immunosorbent assay (ELISA)-based colorimetric reverse transcriptase (RT) assay (Roche Applied Science, Mannheim, Germany).

### Cell Culture

HEK-293T cells were seeded at a density of 35,000 cells per well in 48-well plates using antibiotic-free Dulbecco’s Modified Eagle Medium (DMEM) supplemented with 10 % fetal bovine serum (FBS) and 1 % L-glutamine (Thermo Fisher Scientific), to achieve approximately 70 % confluency. To enhance transduction efficiency of HIV-1 and HIV-2 pseudovirions, cells were first treated with 8 μg/mL polybrene and incubated at 37 °C for 10 minutes. Subsequently, the cells were transduced with 4 ng (reverse transcriptase [RT] equivalent) of HIV-1 or HIV-2, incubated at 37 °C, and collected at 20, 24, and 27 hours post-transduction for analysis by Western blot and real-time qPCR using the Cells-to-CT 1-Step TaqMan Kit ((Thermo Fisher Scientific, Waltham, Massachusetts, USA).

Jurkat cells (ATCC, Manassas, VA, USA) were plated at a density of 70,000 cells per well in 48-well plates containing RPMI-1640 medium (Thermo Fisher Scientific) supplemented with 10 % FBS and 1 % L-glutamine. One hour after seeding, polybrene was added to the culture medium at a final concentration of 8 μg/mL, and the cells were incubated for 10 minutes at 37 °C. Cells were then transduced with 4 ng (RT equivalent) of HIV-1 or HIV-2 pseudovirions. Following transduction, cells were centrifuged using an Eppendorf Concentrator Plus (Sigma-Aldrich, St. Louisan, Missori, USA) for 15 minutes, then incubated for 40 minutes at 37 °C. After incubation, plates were gently shaken to evenly distribute the cells. Samples were collected at 20, 27, 36, 40, and 48 hours post-transduction and analyzed by Western blot and real-time qPCR using the Cells-to-CT 1-Step TaqMan Kit.

### Western blot for the analysis of TFRC

Western blot analysis was performed to detect TFRC expression in HEK-293T and Jurkat cells. Non-treated (naive cells) were used as cell control. At the 27 hour time-point following transduction of HEK293Tcells, the culture medium was removed, and the cells were collected in 200 μL of PBS and centrifuged at 1000 rpm for 4 minutes at 4 °C. The supernatant was discarded, and the cell pellets were lysed using a lysis buffer composed of 50 mM Tris-HCl (pH 7.4), 250 mM NaCl, 5 mM EDTA, 5 mM sodium fluoride, and 0.5% Nonidet P-40. In regards to Jurkat cells, the cells were harvested at 36 and 48 hours post-transduction and processed identically to the HEK-293T cells. Lysates were incubated on ice for 30 minutes, with vortexing for 5 seconds every 10 minutes. The samples were then sonicated using an ultrasonic cleaner (Realsonic) for 2 minutes with 20 second intervals on ice, followed by centrifugation at 14,000 × g for 30 minutes at 4 °C. The supernatant was then transferred to new Eppendorf tubes, and protein concentration was determined using the Pierce BCA Protein Assay Kit (Thermo Scientific, Waltham, MA, USA). Protein samples (15 µg for HEK-293T and 9 µg for Jurkat cells) were analyzed by running on SDS-PAGE (sodium dodecyl sulfate polyacrylamide gel electrophoresis) using a 12% SDS-acrylamide gel. Both the PageRuler Prestained Protein Ladder (10–180 kDa, Catalog No. 26616) and the PageRuler™ Plus Prestained Protein Ladder (10–250 kDa) were used as molecular weight markers. Proteins were transferred to nitrocellulose membrane (Biorad, Hercules, CA, USA), which was then incubated overnight at 4 °C with the primary antibody human CD71 (clone OKT9, eBioscience/Invitrogen, Cat. No. 14-0719-82). The antibody was diluted 1:1500 for HEK-293T samples and 1:2000 for Jurkat samples. To normalize protein loading, a monoclonal anti-β-actin antibody (Sigma-Aldrich) was used at a 1:1000 dilution. Blots were imaged using the Azure Biosystems 600 imaging system following quick incubation with Thermo Scientific SuperSignal ELISA Femto Maximum Sensitivity Substrate (REF 37075). Anti-mouse (Sigma-Aldrich, St. Louis, MO, USA) and anti-rabbit (Biorad, Hercules, CA, USA) were used as secondary antibodies.

### Quantitative Real-time qPCR measurements for the detection of TFRC

RNA expression analysis was conducted to assess TFRC expression using the Cells-to-CT 1-Step TaqMan Kit (ThermoFisher Scientific) following the manufacturer’s protocol. The culture medium was removed from both control (native) and HIV-1 and 2 transduced cells, and then the cells were collected in 200 μL of PBS. Samples were centrifuged at 1000 rpm for 4 minutes at 4 °C. After centrifugation, the supernatant was discarded, and the cells were lysed in 49.5 μL of room-temperature lysis solution supplemented with 0.5 μL DNase for 5 minutes, followed by the addition of 5 μL of room-temperature stop solution and the samples were incubated for 2 minutes. Each qRT-PCR reaction included the following: 2 μL of cell lysate, 5 μL of TaqMan 1-Step qRT-PCR Mix, 1 μL of TaqMan Gene Expression Assay (20X), and 12 Nuclease-Free Water to a final volume of 20 μL. Reactions on ice were loaded onto an Axygen 384-well PCR Microplate (Axygen, nion City, California, USA) and run on a LightCycler 480 Instrument II (Roche Holding AG, Basel, Switzerland), following the Standard cycling conditions provided in the kit manual.

### Transfection of HEK-293T cells with Y44A inactivated Tat construct

HEK-293T cells (250,000 cells per well) were seeded in a 24-well plate using antibiotic-free DMEM supplemented with 10 % FBS and 1 % L-glutamine. Once the cells reached ∼70% confluency, the cells were transfected with 500 ng of either the packaging plasmid psPAX2, CGP (HIV-2 packaging plasmid) or Y44A (HIV-2 CGP packaging plasmid containing a deactivating mutation in the Tat protein). Y44A was previously constructed [40] by introducing the inactivating mutation into the HIV-2 *tat* gene within the HIV-2 CGP vector.

Lipofectamine 3000 LTX reagent and Opti-MEM (Thermo Fisher Scientific) were used for the transfection, following the manufacturer’s protocol. Briefly, 500 ng of plasmid DNA was incubated with 1 µL LTX3000 Reagent in a total volume of 50 µL OPTI-MEM. During the 20-minute incubation period, the medium was removed from the cells and replaced with transfection medium, with a total volume of 500 µL per well (complemented with DMEM). The culture medium was replaced 5 hours post-transfection, and the cells were subsequently incubated and collected by pipetting 27 hours post-transfection in 500 μL of PBS. The samples were centrifuged at 1000 rpm for 4 minutes at 4 °C, and the supernatant was discarded. The resulting cell pellet was used for Western blot analysis to detect TFRC expression.

### Measurement of total iron content in transduced Jurkat cells

To determine the iron content in transduced Jurkat cells, approximately 1 × 10^6^ cells were seeded in a 6-well plate. After one hour incubation, the cells were transduced with 50 ng (RT equivalent) of HIV-1 or HIV-2 pseudovirions. Two hours post-transduction, Bovine (Holo form) Transferrin (Thermo Fisher Scientific) was added to the medium at a final concentration of 50 µg/mL. Cells were incubated at 37 °C and collected 48 hours post-transduction. The collected cells were transferred to 15 mL tubes and centrifuged at 870 rpm for 5 minutes at 24 °C. The supernatant was discarded, and the cell pellets were re-suspended in 2000 µL of cold PBS. Iron content was measured using the Cell Total Iron Colorimetric Assay Kit (Invitrogen, Waltham, Massachusetts, USA), following the manufacturer’s protocol. A total of 1 × 10^6^ cells were counted and re-suspended in 400 µL of 0.9 % NaCl, then centrifuged at 300 × g for 10 minutes at 4 °C. After discarding the supernatant, the cell pellet was homogenized in 200 µL of the assay buffer and incubated on ice for 10 minutes. The lysate was then centrifuged at 15,000 × g for 10 minutes, and the resulting supernatant was collected for iron quantification. Statistical analysis was performed according to the manufacturer’s instructions.

## Results

### Induction of TFRC expression by HIV-2 in HEK-293T cells

To investigate the regulation of TFRC expression by HIV-1 and 2, HEK-293T cells were transduced with HIV-1 and HIV-2 pseudovirions, and samples were collected at 20, 24, and 27 hours post-transduction. Using qPCR to analyse the expression of TFRC, in HEK-293T cells, we could see a significant upregulation of TFRC at 27 hours in cells transduced with HIV-2, compared to those transduced with HIV-1 (Figure 1A). TFRC protein expression was also assessed by Western blotting of cell lysates at the 27 hours’ time-point (Figure 1B).

**Figure 1.**
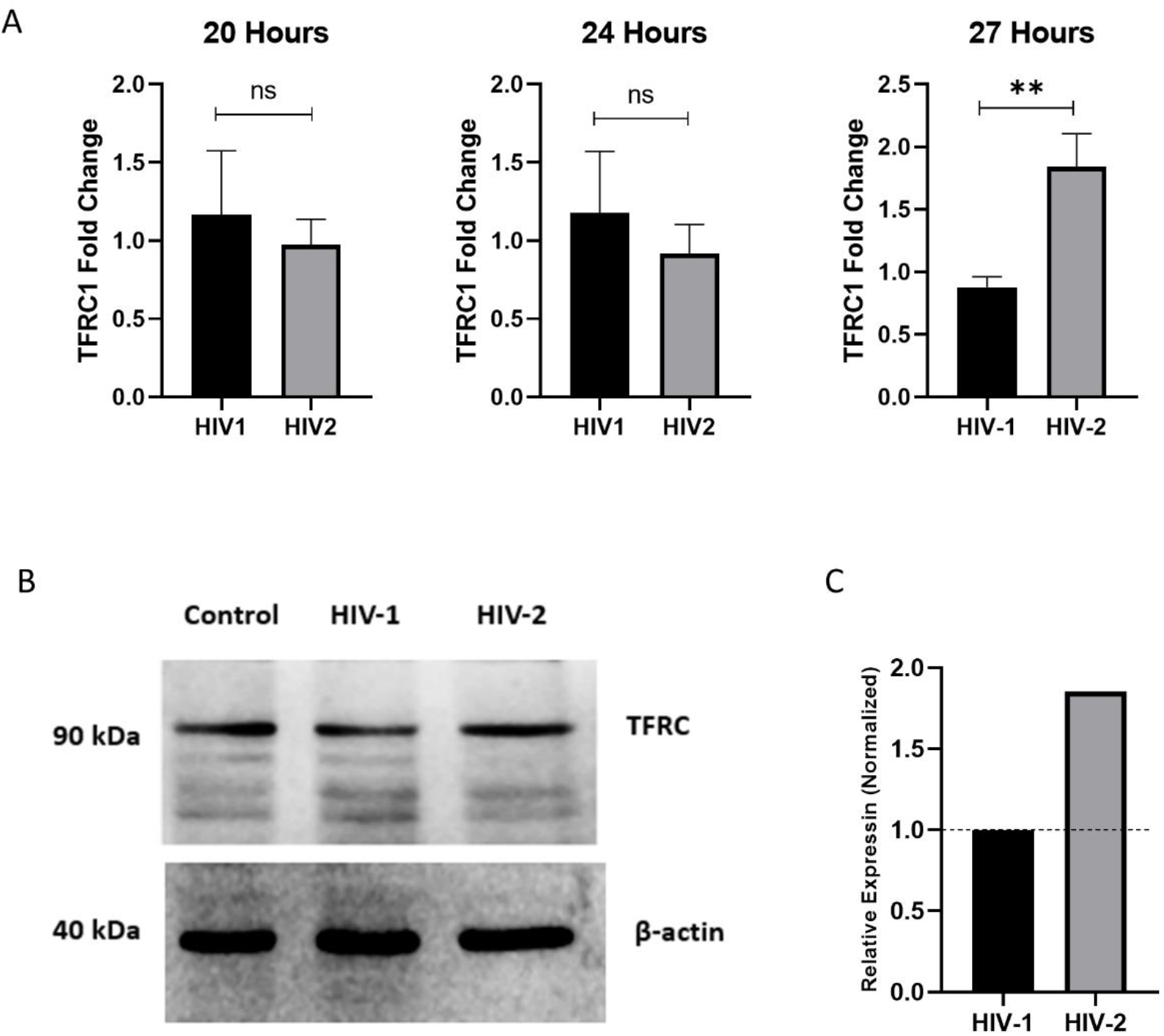
**A)** Real-time qPCR analysis of transduced HEK-293T cells with HIV-1 and HIV-2 pseudovirions. TFRC fold-change is shown after 20hr, 24hr, and 27hr post-transduction. Results from triplicate measurements show significant upregulation of TFRC at the 27hr time-point, compared to HIV-1. The statistical analysis was performed by Graphpad prism 8.0.1. **B)** Representative Western blot analysis of TFRC at 27 hours post-transduction in HIV-1 and HIV-2 transduced cells, from at least 3 independent experiments. Levels of TFRC in HIV-2 transduced cells were higher than those in HIV-1 transduced cells as detected by densitometric analysis (C). Relative expression was compared to HIV-1-transfected cells.

### Induction of TFRC expression by HIV-2 in Jurkat cells

In transduced Jurkat cells, samples were collected at 20, 27, 36, 40, and 48 hours post-transduction. Untreated cells were used as controls. Using GAPDH for normalization, fold change of TFRC transcripts were compared between HIV-1 and HIV-2 transduced cells.

Compared to HEK-293T cells, the upregulation of TFRC was observed at 36 hours post-transduction in the case of HIV-2 transduced cells, an upregulation that remained significant at 36 and even 40 hours post-transduction. At 48 hours post-transduction, there was no significant change in TFRC regulation between the two pseudoviruses (Figure 2A). Protein expression was assessed by Western blotting the cell lysates. Despite the upregulation of TFRC transcripts by HIV-2 in the 36 hour time-point, this was not reflected on the protein level, as levels of TFRC were lower compared to that found in cells transduced with HIV-1 and the cell control. This downregulation on the protein level was also visible at the 48 hour time-point.

**Figure 2.**
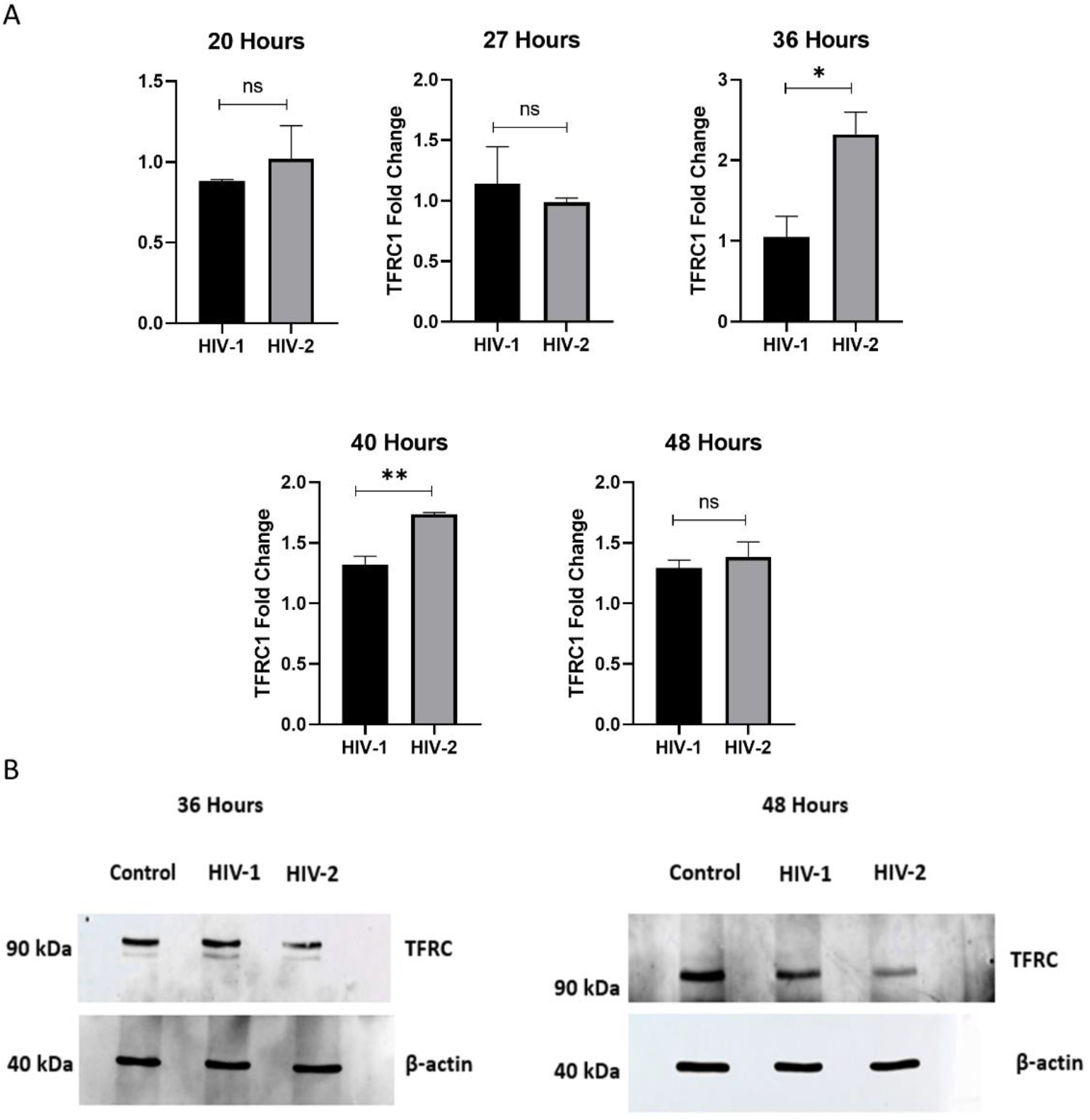
**A)** Real-time qPCR analysis of HIV-1 and HIV-2 transduced Jurkat cells. TFRC fold-change is shown after 20, 27, 36, 40, and 48 hours post-transduction. Significant upregulation of TFRC was observed after the 36 hour time-point, persisting until 40 hours post-transduction with HIV-2. **B)** Western blot analysis of TFRC expression in Jurkat cells transduced with HIV-1 and HIV-2 at 36 hours and 48 hours post-transduction. Although the TFRC transcripts were significantly upregulated by HIV-2 at the 36 hour time-point, we observed a downregulation at the protein level. Also, at 48 hours post-transduction, the level of TFRC protein remained lower than that of the control and HIV-1 transduced cells. Results are representative of at least 3 independent experiments.

### Modulation of TFRC by HIV-2 is Tat dependent

To investigate the culprit behind the modulation of TFRC by HIV-2, HEK-293T cells were transfected with an HIV-2 packaging plasmid encoding an inactive Tat mutant (A44Y). As controls, cells were transfected with either a wild-type HIV-2 packaging plasmid or the psPAX2 HIV-1 packaging plasmid. Quantitative analysis revealed that TFRC expression was reduced in cells transfected with the HIV-2 Tat-defective plasmid compared to those transfected with the wild-type HIV-2 construct. Moreover, TFRC levels were higher in cells expressing wild-type HIV-2 compared to those transfected with the HIV-1 packaging plasmid, indicating that functional HIV-2 Tat contributes to the modulation of TFRC expression (Figure 3).

**Figure 3.**
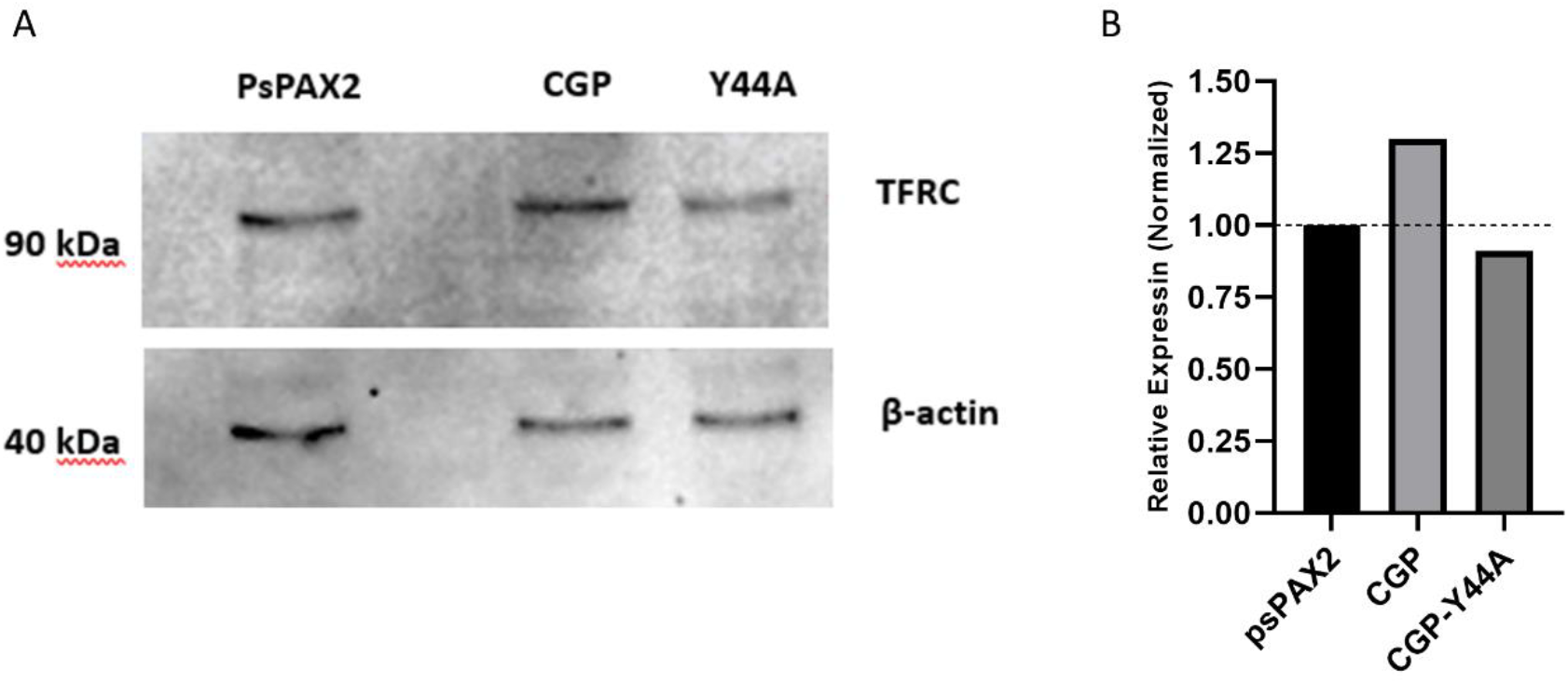
**A)** Western blot of HEK-293T cells transfected with HIV-2 packaging plasmid coding for wild-type (CGP) and mutant (CGP-Y44A) Tat protein. TFRC level was lower in cells transfected with the inactivated Tat mutant. B) Densitometric analysis of TFRC bands detected by western blot. Relative expression was calculated compared to the HIV-1-based, psPAX2-transfected cells.

### Assessing the impact TFRC modulation by HIV-2 on cellular iron uptake

We determined the total iron concentration in Jurkat cells, 48 hours after transduction with HIV-2 using the Cell Total Iron Colorimetric Assay Kit. The results indicated that Jurkat cells transduced with HIV-2 exhibited higher intracellular iron content compared to those transduced with HIV-1 and the control native cells (Figure 4).

**Figure 4.**
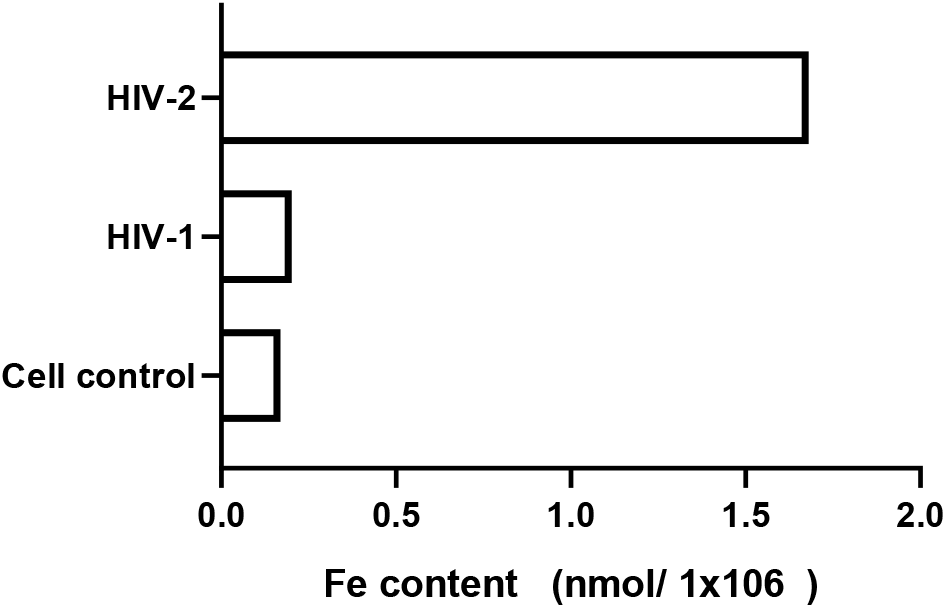
Intracellular iron content in Jurkat cells transduced with HIV-1 and HIV-2 pseudovirions. Jurkat cells transduced with HIV-2 pseudovirions exhibited higher intracellular iron levels compared to those transduced with HIV-1. Control cells are non-transduced cells.

## Discussion

The findings of this study provide novel insights into the differential regulation of TFRC expression by HIV, highlighting the role of HIV-2 Tat in its modulation, and the potential implications for viral entry and intracellular iron homeostasis. Our results demonstrate that transduction of HEK-293T cells with HIV-2 pseudovirions results in a significant upregulation of TFRC mRNA at 27 hours post-transduction compared to HIV-1-transduced cells, as measured by qPCR. This transcriptional upregulation was corroborated at the protein level accessed by western blotting, indicating that HIV-2, and not HIV-1, specifically enhances TFRC expression in this cell type. On the other hand, Jurkat cells exhibited a delayed but statistically significant increase in TFRC transcript levels at 36 and 40 hours post-transduction with HIV-2, a pattern not observed in HIV-1-transduced cells. These temporal and cell-type-specific patterns underscore the distinct molecular interactions between HIV-2 and the host cells. Notably, the delayed induction of TFRC expression in Jurkat cells suggests that HIV-2 may engage different regulatory pathways depending on the cellular context.

Despite the upregulation of TFRC mRNA in HIV-2-transduced Jurkat cells, this effect was not mirrored at the protein level. TFRC protein abundance was reduced compared to both HIV-1-transduced and control cells at 36 and 48 hours post-transduction. This discrepancy suggests the involvement of post-transcriptional mechanisms such as translational inhibition or enhanced proteasomal degradation that selectively impair TFRC protein expression in HIV-2-transduced Jurkat cells [42-45]. Mechanisms including microRNA-mediated translational repression or ubiquitin-proteasome system targeting have been implicated in HIV-related host protein modulation and could be implicated in the modulation of TFRC either directly or indirectly [46,47].

To further analyze the molecular basis of HIV-2–mediated TFRC regulation, we employed an HIV-2 packaging construct encoding an inactive HIV-2 Tat mutant (A44Y). Transfection of HEK-293T cells with this mutant led to a significant reduction in TFRC expression relative to cells expressing wild-type HIV-2 Tat, implicating functional Tat as a key modulator of TFRC upregulation. Furthermore, the finding that the wild-type HIV-2 packaging plasmid induces higher TFRC expression than its HIV-1 counterpart does supports the hypothesis that HIV-2 Tat may engage the host transcriptional machinery through a mechanistically distinct pathway compared to HIV-1. Chromatin immunoprecipitation and promoter-binding assays could provide further insight into the mechanism by which TFRC expression is modulated, and their contribution to iron metabolism or viral entry.

Functionally, HIV-2-mediated TFRC modulation detected by elevated mRNA level but decreased protein level corresponded with a dramatic increase in intracellular iron content in Jurkat cells, as measured at 48 hours post-transduction using a total iron colorimetric assay suggesting an overall elevated iron import into the cells. On the contrary, HIV-1 transduction did not alter the intracellular iron level indicating that only HIV-2 may influence iron uptake and cellular iron homeostasis. Elevated intracellular iron levels have been associated with increased oxidative stress and immunopathology during HIV infection [48]. Comprehensive profiling of additional iron-regulatory genes and proteins would help elucidate the broader impact of HIV-2 on iron metabolism.

The distinct regulatory patterns observed in HEK-293T and Jurkat cells emphasize the importance of cellular context in host–virus interactions. The earlier TFRC upregulation in HEK-293T cells compared to Jurkat cells may reflect differences in viral entry kinetics, transcriptional landscapes, or availability of host cofactors [49-51]. The lack of significant TFRC influence by HIV-1 in either cell type further highlights the unique capabilities of HIV-2 to manipulate entry pathways. These results contribute to the growing body of evidence indicating that HIV-1 and HIV-2, despite their genetic similarities, exhibit distinct molecular strategies for host modulation [7,50,51].

A major limitation of our study is that we were unable to monitor the modulation of TFRC beyond the early phase of viral transduction, due to the use of self-inactivating, replication-incompetent pseudovirions. This constraint also prevented us from assessing whether such modulation influences viral entry. Furthermore, since we employed VSV-G–pseudotyped virions, we were unable to independently evaluate the potential enhancement of viral entry resulting from upregulation of TFRC. Nevertheless, considering the broad receptor usage and tissue tropism characteristic of HIV-2, the possibility that TFRC may serve as a viable and advantageous entry factor warrants further investigation. Such a mechanism could potentially circumvent the well-documented phenomenon of superinfection resistance observed in most retroviruses [52]. Notably, HIV-2 is known to exhibit weaker downregulation of CD4 and lacks the Vpu protein found in HIV-1, factors that may contribute to a more permissive environment for superinfection [53,54].

In conclusion, our study demonstrates that HIV-2, probably through the activity of its Tat protein, modulates TFRC expression in a cell type– and time-dependent manner, leading to increased intracellular iron accumulation. These findings reveal a novel facet of HIV-2–host interactions that may impact viral entry and replication dynamics, oxidative stress responses, and immune modulation. Elucidating the mechanisms by which Tat regulates TFRC, as well as the downstream consequences of this modulation, will provide valuable insights into the differential pathogenesis of HIV-1 and HIV-2, and may uncover new targets for therapeutic intervention.

## Acknowledgments

We are grateful to Joseph P. Dougherty from the Robert Wood Johnson Medical School, NJ, USA, for providing us with the HIV-2 CGP vector. The authors would also like to extend their gratitude to the staff of the Laboratory of Retroviral Biochemistry for their continued support.

## Author Contributions

A.S.A: Performed experiments, collected and analysed data, drafted the manuscript, IWK: Assisted in the production of pseudovirions, M.M: designed the experiments and interpreted the results, wrote the manuscript; J.T: supervised and funded the project, reviewed and modified the manuscript. All authors have read and agreed to the published version of the manuscript.

